# Wavelet analysis of human recombination rates demonstrates divergence on fine scales

**DOI:** 10.64898/2026.06.11.731649

**Authors:** Clare Horscroft, Andrew Collins, Reuben J Pengelly

## Abstract

**Background:** Recombination rates can be estimated across the genome, underpinning genetic analyses such as identification of regions under selection. Accurate recombination mapping requires observing a large number of recombination events, necessitating large sample sizes to achieve high resolution. This can be prohibitive to some analyses, so population-based estimates can also be used, leveraging the increasing population genomic data available to researchers.

**Objective:** This study aimed to determine the extent to which population-based recombination maps from different human populations are similar and assess to what extent they can be used interchangeably.

**Methods:** Wavelet analysis was employed to decompose recombination rate signals along a chromosome and evaluate the proportion of variance explained at different scales. This method also enabled the assessment of correlations across scales and identification of regions with high coherence between datasets. The analysis focused on a region of chromosome 22 in human populations of European and African ancestry.

**Results:** Recombination rates are not closely conserved across populations, with the greatest divergence observed at fine scales. Coherence between populations varied significantly across all scales.

**Conclusion:** As recombination maps differ substantially between human populations, for genetic analyses involving recombination maps, it is recommended to use maps specific to the population under study.

## INTRODUCTION

The study of evolution and natural selection from a genomic perspective continues to increase our understanding of the genetic and phenotypic variation observable within and between human populations. This is especially pertinent in medicine where there are implications in many areas including disease susceptibility, immunology, and drug metabolism. Many methods for identifying evidence of natural selection in the genome rely on the statistical relationships between alleles [1, 2]. The process of recombination in meiosis breaks down these relationships, and so it is important to account for the effects of recombination when applying these methods [3].

Meiotic recombination is the process in which homologous chromosomes crossover and exchange genetic material to make a new chromosome containing genetic information from both the original chromosomes. There are two main evolutionary benefits to recombination: firstly it is a mechanism in which multiple beneficial mutations, which exist on different haplotypes, may be brought together with potential fitness benefits; and secondly, it may reduce the accumulation of deleterious mutations (reducing the impact of Muller’s ratchet) [4, 5]. Recombination rates are not uniform across the genome: there are observable hotspots where recombination is much more frequent and cold spots where little recombination takes place. Recombination is also positively correlated with distance from chromosome centromeres, with little recombination taking place near the centromere [6].

Recombination hotspots are strongly linked with the activity of the zinc-finger protein PRDM9 [7–9]. The zinc-finger of PRDM9 binds with particular DNA motifs and contains a SET domain which catalyses the trimethylation of lysine 4 of histone H3 (H3K4me3) [10], and to a lesser extent, lysine 36 (H3K36me3) [11]. Regions of the genome enriched for H3K4me3 and H3K36me3 have been found to be more likely to recombine, hence the correlation between the PRDM9 DNA binding motifs and recombination hotspots [12–14]. Humans and chimpanzees (*Pan troglodytes*) share very few hotspots regardless of the similarity of the genetic code, due to the rapid evolution of PRDM9 causing different DNA binding motifs to be preferred [15]. Globally, within human populations there are different variants of PRDM9 that cause hotspots to arise in different places along the genome, with the frequencies of these variants differing between populations [16].

When considering the relationships between loci, there are two main approaches through which this can be assessed, linkage and linkage disequilibrium (LD). To define explicitly, linkage is the phenomenon whereby loci on the same chromosome do not segregate independently during meiosis, given that they will co-segregate in the absence of intervening recombination events [17]. Linkage maps are based on family data in which recombination events can be directly inferred between individuals including mother-father-child trios [18] with more information coming from families with multiple children, and blastocysts derived through *in vitro* fertilisation can also be utilised to increase power [19, 20]. Linkage maps have the benefit of allowing sex-specific analysis; however, they are low resolution and can only show recombination events between genotyped individuals over a small number of recent generations.

In contrast to linkage, LD is the statistical association of alleles between loci, quantified on a population basis [17]. LD-based recombination maps use population genetic data to estimate recombination patterns across the genome, for example as reported by Myers et al. [21]. By using population data, the pattern of recombination events over many historical generations may be captured, and thus the resolution is much higher compared to conventional linkage analysis. Furthermore, the attainable resolution increases with the ever-growing availability of whole genome sequencing (WGS) data [22]. However, comparisons of these LD-based recombination maps with linkage maps show that the residual impacts of additional processes such as selection, genetic drift and population bottlenecks may distort the maps to some degree, confounding true recombination estimates [23, 24]. The authors of the LDhat software for estimating recombination rates from LD patterns acknowledge that it is difficult to obtain the pure recombination rates given these confounding factors [25, 26]. Reed and Tishkoff [27] showed that some software for estimating recombination rates from LD can be confounded by positive selective sweeps, causing false-positive hotspot detection. It has been argued that selective sweeps should not affect the coalescent-based method used by LDhat and, at worst, would produce a small decrease in recombination estimates around a strong sweep [28, 29]. However, simulations performed by Chan et al. [30] show some evidence for false positive hotspot detection around positive sweeps using LDhat.

Detecting positive selection sweeps located near a recombination hotspot is challenging. Generally, sweeps cause a lengthening of haplotypes, distortions in the allele frequency spectrum, and an increase in the association between alleles [31]. Frequent recombination over generations results in shortening of haplotypes and relationships between alleles decline quickly, masking sweep signatures. Conversely, areas of the genome with very low recombination rates may be mistaken for a sweep, as the correlation between alleles would be higher than elsewhere in the genome. Therefore, correcting or adjusting for recombination rates, where known, would be advantageous [32].

Different populations within a species can have different recombination hotspots [30, 33, 34]. However, the overall pattern of recombination may remain similar on a macro scale, for example the correlation with distance from the centromere [32]. This raises the question of the validity of using a single generic recombination map for a species for all populations within that species, and specifically, at what resolution that would be appropriate, if at all.

Wavelet transforms are mathematical methods related to Fourier analysis developed for processing signals [35]. While traditionally used for time series data, wavelets can also be applied to any phenomena that exist along a continuum. Wavelet analysis extracts information about the signal at different scales, from very short-term changes to overarching long-term trends. Wavelets work similarly to a moving average, but without having to choose a fixed window size. Wavelets have proven to be incredibly useful for signal compression and smoothing and have been widely used for image and sound processing [36–38].

There are many examples of wavelet techniques being applied in a range of genomic applications, including the detection of copy number variants [39–41], microarray de-noising [42], and downsizing of genetic data for more efficient analysis [43]. Wavelets have been used for analysing signals, for example in analysing the features found around indels [44], features of admixture in human populations [45], and dependencies between different genetic motifs [46]. Most relevant to this study are analyses involving recombination rates. Bherer et al. [47] used wavelets to investigate sexual dimorphism in recombination rate, finding that male and female rates differed most at the fine scale, and that the power spectrum for both rates showed the majority of variance in recombination rates was found at the mid-scale between 16 and 64 kb. Finally, Chan et al. [30] used wavelet analysis to assess the difference in recombination rates between two populations of *Drosophila melanogaster*, reporting that correlations between the populations increased with scale.

The application of wavelets to genomic data allows for the analysis of signals at different scales along a chromosome. One of the attractive things about wavelet analysis is the ability to look at fine scale and wider scales simultaneously, without having to choose arbitrary window sizes that could skew results, particularly because recombination hotspot estimation has been shown to be sensitive to window size [48]. The contribution of each scale to the overall variance of the signal can be ascertained, and this is known as the power spectrum. This allows for easy comparison between multiple datasets across different scales.

We evaluate here wavelet analysis of recombination patterns in four different populations and compare the patterns of variation along a typical chromosome to ascertain to what extent recombination structure is conserved on different scales. This provides insights into what extent recombination maps can be used interchangeably in studies of different populations.

## MATERIALS AND METHODS

### Data

Four population WGS datasets were considered (Table 1). The two European descent datasets (CEU and Wellderly) and two African datasets (Baganda and Zulu) were chosen for analysis to allow comparison between African and European genomes whilst also comparing within the continental groups to benchmark the expected variation. Variant call data based on reference genome GRCh37 (hg19) was utilised, for all datasets.

**Table 1.** | Samples used for analysis.

| **Table 1 \| Samples used for analysis** |  |  |
| --- | --- | --- |
| **Population label** | **Ancestry** | **Source** |
| Welllderly | European descent | Erikson *et al*, 2016 [49] |
| CEU | European descent | 1000 Genomes Psroject, 2015 [50] |
| Baganda | Ugandan | Gurdasani *et al* (2014) [51] |
| Zulu | South African | Gurdasani *et al* (2014) [51] |

In the Wellderly dataset, principal component analysis (PCA) was used to identify a homogenous population (Supplementary Figure 1) which was used for further analysis. The CEU data refer to Utah residents with Northern and Western European ancestry from the 1000 Genomes Project [50]. The Baganda and Zulu datasets are from the African Genome Variation Project [51] and contain individuals from Uganda and South Africa respectively), Chromosome 22 was utilised for analysis as a representative human autosomal chromosome for computational feasibility. The datasets were cleaned using PLINK [52] by removing SNPs with MAF ≤ 0.05, missing data ≥ 10%, or Hardy-Weinberg equilibrium p-value ≤ 0.001. For all subsequent analyses, 96 samples were randomly selected where more were present, to ensure parity between datasets and to meet software limitations for data size.

### Estimating recombination rates

Recombination rates were estimated using the LDhat software [25]. The premade lookup table with parameters *n* = 192 and θ = 0.001 was used. The convert program within LDhat was used to randomly select 96 samples from each dataset and remove SNPs with ≥ 10% missing values. The number of iterations was set to 10,000,000 with sampling every 5,000 iterations and a block penalty of 5. The first 100,000 iterations were discarded.

### Wavelets

All wavelet analysis was undertaken using the R programming software version 3.3.0 [53]. The R packages used were “intervals” version 0.15.1 [54], “wmtsa” version 2.0-3 [55] and “biwavelet” version 0.20.17 [56].

The datasets were trimmed to the chromosomal region chr22:20000428-51218377, which is entirely euchromatic. Following Booker *et al.* [57] the data were converted from ρ = 4N_e_r per kilobase (Kb) into centimorgans per megabase (cM/Mb) using a map length of 60.67105 cM calculated from the Bherer *et al.* [47] paper. Note that there is no reason to believe that chromosomes in African populations recombine more or less than Europeans (just in different places) and indeed the African-Enriched recombination map produced by Hinch *et al.* [33] resulted in a very similar map length of 60.2716 cM over the same region, so the Bherer figure was used for all datasets. The ρ values were cumulatively summed, weighting for the distance between SNPs, and then proportioned based on the map length to get cumulative cM. This was then converted to cM/Mb based on the distances. The data were then binned into 1 Kb bins with interpolation using the code provided by Bherer *et al.* [47], excluding any gaps bigger than 50 Kb. The datasets were merged and the longest section of chromosome 22 where all four datasets had complete data was utilised for downstream analysis to avoid differential data quality interfering with analyses. Finally, the region chr22:29527000-45911000 was selected to be analysed as it is at the centre of the region and contains a number of bins equal to a power of two (16,384 bins = 2^14^) as discrete wavelet transforms work by decomposing the data into a number of scales *j* where the signal contains 2^j^ data points. Constraining the data this way avoids having to pad the data artificially or use other fixes which can have drawbacks [35].

Maximal overlap discrete wavelet transforms (MODWTs) were undertaken using the Haar wavelet, and continuous wavelet transforms (CWTs) utilised the Morlet wavelet with parameter ω_0_ = 6 as in Chan *et al.* [30] due to its wide use and appropriateness for extracting features from signals [58].

## RESULTS

### Recombination maps

Cumulative inferred centimorgan (cM) maps were created for each of the four WGS datasets using the LDhat software [25], (Supplementary Figure 2). Maps are flatter where there is less recombination and sharp increases in slope correspond to regions of increased recombination intensity. The maps for the four populations follow the same macro pattern. The CEU map shows a sharper increase in slope around 22 Mb from the origin and both European maps show a sharp increase around 40 Mb which is not seen in the African maps, suggesting a recombination hotspot which is not shared between the populations.

To validate the use of the LDhat software as an estimator for recombination rates, the CEU results were compared to the refined European sex-averaged linkage map created by Bherer *et al.* [47] for chromosome 22, with a τ coefficient of 0.41 at a 1 Kb scale (p < 2.2×10^16^; Supplementary Figures 2 & 3). The LDhat model was thus deemed a reasonable approximation of a linkage map. The LDhat map contained evidence of now extinct hotspots that are not present in linkage maps generated from data containing only observed meioses. Other differences could be due to other confounders to the LDhat model such as selection, drift, and demographic changes.

### Power spectrum

The power spectrum can be calculated from the detail coefficients of a discrete wavelet transform (DWT). Detail coefficients were calculated on a range of scales, from 2 Kb through to ∼16 Mb. The properties of wavelets allow ascertainment of what proportion of the variance from the original signal is attributable to each scale, and only that scale.

Figure 1 shows the power spectrum of the log_10_ transformed European and African datasets. There is a clear peak within the 16-64 Kb range. This may reflect the expectation that there is, on average, around one recombination hotspot every 50 Kb [21].

**Figure 1.**
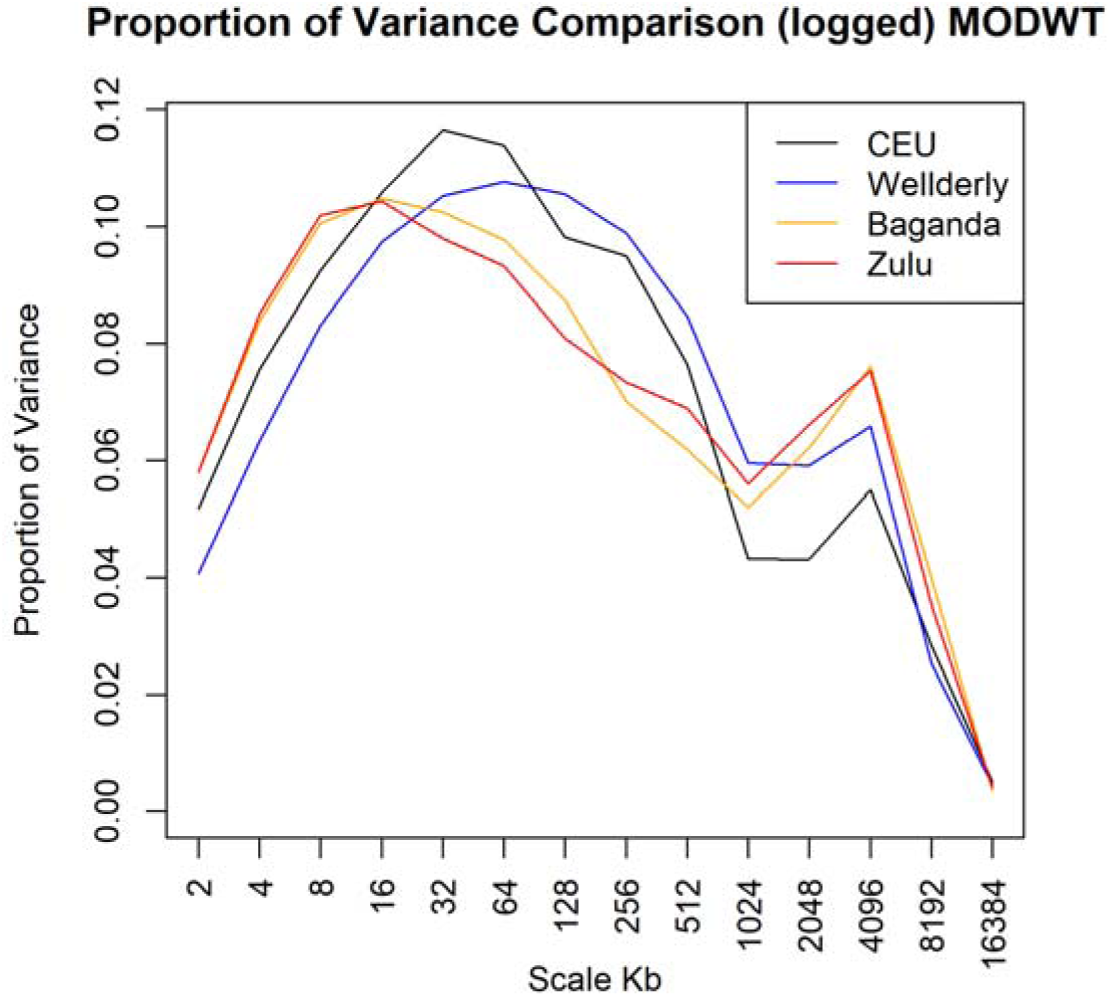
Proportion of variance captures at varying scales. Maximal Overlap Discrete Wavelet Transform (MODWT) power spectrum for the four datasets CEU, Wellderly, Baganda and Zulu, for a 16 Mb region of chromosome 22. The graph shows the proportion of variance contained within each scale for the log_10_ signals.

### Correlations

The wavelet decomposition allows for the correlations between the detail coefficients to be calculated between datasets at each scale. Detail coefficients describe the change in the signal. Correlation between detail coefficients therefore implies that as one signal changes (at that scale), as does the other. As the data are not normally distributed, Kendall’s Tau was applied. Figure 2 shows the correlations between the detail coefficients for each pair of datasets.

**Figure 2.**
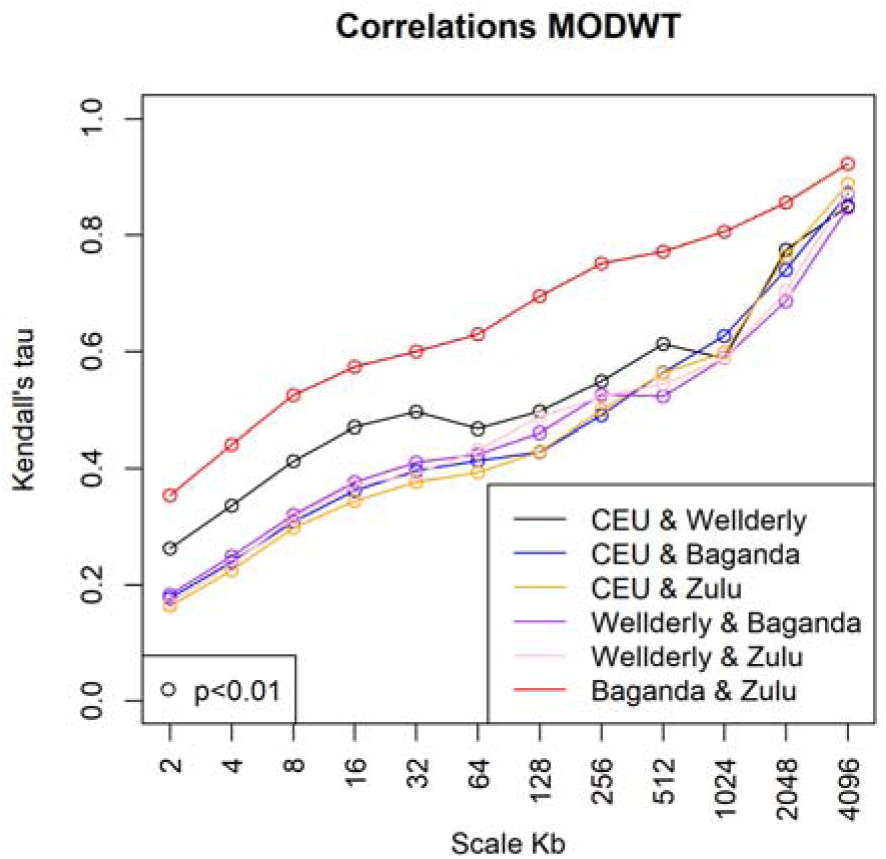
Correlations between datasets at varying scales. The correlations between the detail coefficients of the maximal overlap DWT of the log_10_ transformed datasets. Circles indicate correlations with a p-value less than 0.01.

Figure 2 shows an upward trend for each pairing – that is, the wider the scale, the higher the correlation between the detail coefficients. Changes in the original signal on the narrow scale are much less correlated than changes in the signal on the wider scale. This could be because recombination tends to cluster leaving cold spots in between, where the precise location of hotspots can be mobile over generations [59]. It is established that hotspots can reposition rapidly: humans share very few hotspots with chimpanzees despite the similarity in our DNA sequences [15]. This is part of the so-called “hotspot paradox”, where hotspots are more likely to be sites for a recombination event, however, through the mechanisms of this process and biased gene conversion the site becomes less likely to recombine [60]. The upwards trend in Figure 2 therefore can be explained by the repositioning of hotspots at the fine scale while the overall recombination rate averages out as the scale increases.

The African datasets are more correlated with each other on every scale than any other pairing, which was unexpected. Potential reasons for this will be discussed further later. Using the correlations between the European pair as a baseline, the European and African pairs differ mostly at the fine scale, and as the scales increase the correlations become very similar to what would be expected for two samples from the same (European) population. The figure indicates that the largest differences between the European and African datasets are at the fine scale.

### Continuous Wavelet Transformation

Figure 3 shows the CWT for each of the four datasets, plus their underlying data below. The colours on the graphs represent the magnitude of the detail coefficients – for example if the recombination rate was flat for the whole region, the graph would be entirely blue due to there being no changes across the area at any scale. The black lines indicate areas that are significant at the 5% level compared to a background power spectrum. This is calculated using a χ^2^ test, comparing to a red noise null power spectrum modelled using an autoregressive process with a lag of 1 [61]. The graphs show that the significant power differences are present at the narrowest scales and the widest scales. The narrow scales also contain most of the blue, low power areas. The proportion of variance in the power spectrum for the fine scale was lower than the mid-range scale, despite the abundance of high-power areas, because of these regions of static rates.

**Figure 3.**
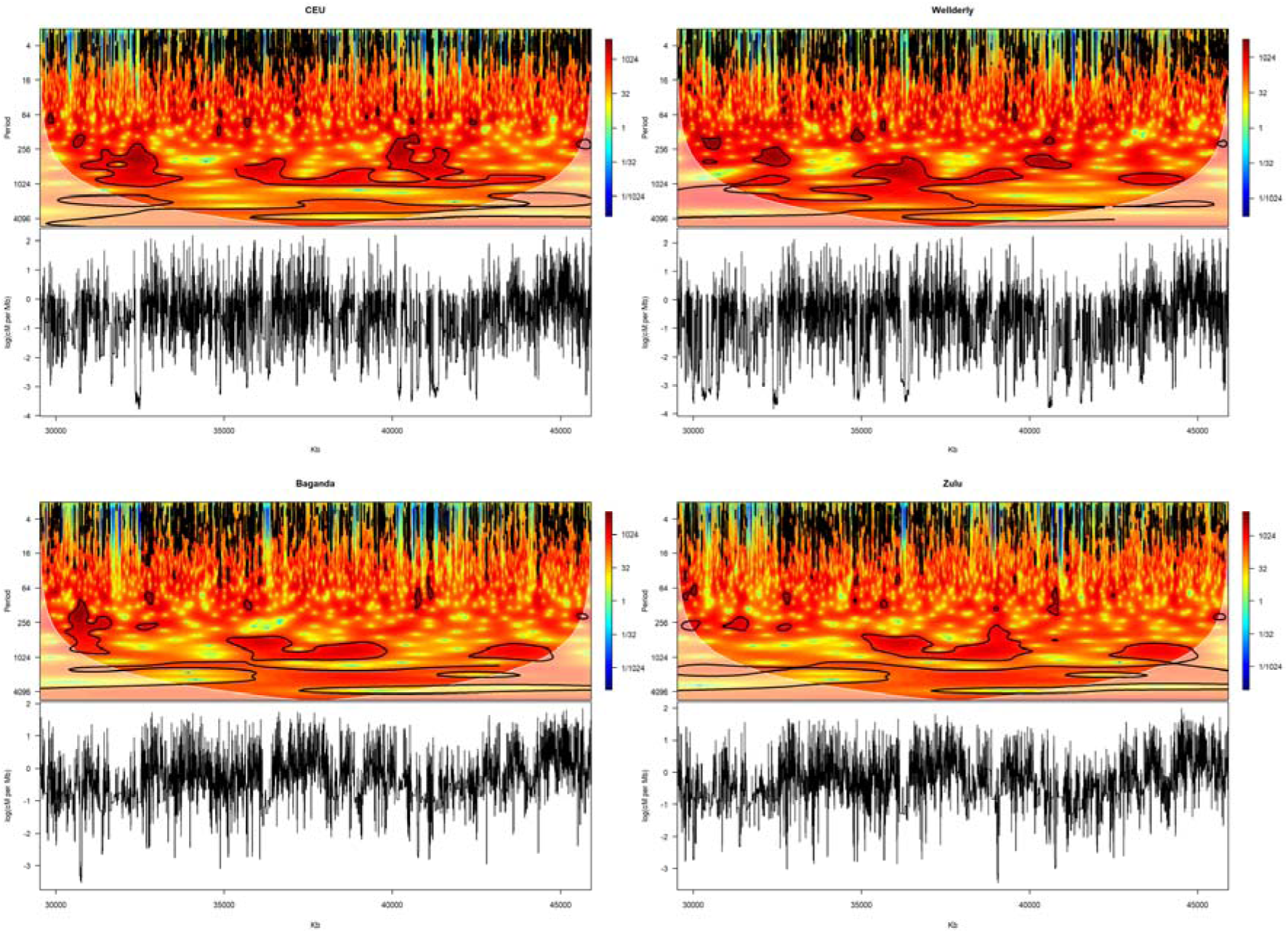
Continuous wavelet transformation. The CWT of the logged datasets for the four populations showing the CWT and the original signal below, with kb position (horizontal) against wavelet period (in kb, vertical). The colours indicate the magnitude of the detail coefficient – a flat, constant recombination rate would be dark blue. The black outlined areas are significant at the 5% level. The white shaded areas are under the cone of influence, where data may be affected by edge effects.

**Figure 4.**
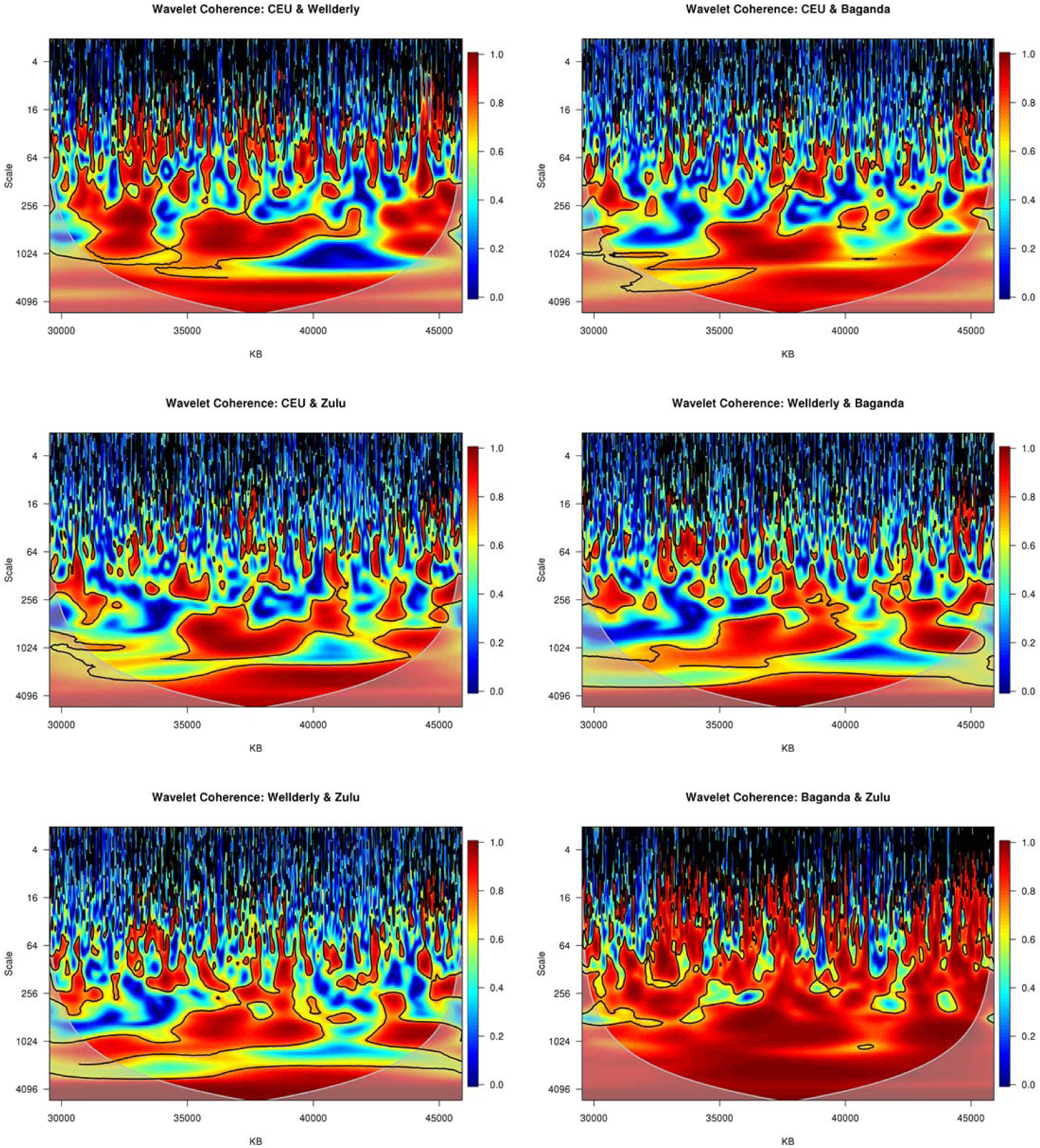
Wavelet coherence analysis showing correlation between populations. Wavelet coherence analysis applied to each pair of recombination rate datasets, with kb position (horizontal) against correlation scale (in kb, vertical). Red colours indicate higher correlation between the datasets at that location and scale. The cone of influence is shown in white. The black lines border regions of the graph at 5% significance. The highly correlated red area on the CEU & Wellderly graph from the 256 to 1024 Kb scale around 32.5 Mb is not replicated in the European/African graphs or indeed the Baganda & Zulu graph. This is due to a roughly 150 Kb wide cold spot present in both European datasets but not in the African data (Supplementary Figure 4).

### Wavelet Coherence

This analysis assesses whether the correlations in Figure 3 are globally representative of the whole region or whether there are differences in the correlation not only for multiple scales but also at different points along the chromosome. It works by calculating a smoothed correlation coefficient as defined in Torrence and Webster [62]. To test for significance, a Monte Carlo method was applied, using 1,000 Monte Carlo randomisations as recommended in Grinsted *et al*. [58]. The results of the wavelet coherence analysis for each pair of datasets can be seen in Figure. The first striking feature is the strong correlation between the Baganda and Zulu data across most of the region, and on medium to wide scales. Despite this strong correlation, there are many pockets of weak/no correlation. This shows that even in these relatively similar populations there are still differences in the changes in the recombination rate at the fine scale. In contrast to this, the graph of the two European datasets (CEU and Wellderly) has many more regions exhibiting weak correlations, especially in the medium scales (64 to 256). The graphs comparing the European and African datasets all show a lack of correlation between the changes in the two signals on scales of around 64 Kb and less, with higher correlation at the wider scales – though noting a high proportion of the red areas are below the cone of influence.

## DISCUSSION

Through wavelet-based comparison of the recombination structure of chromosome 22, it is clear that recombination rates are not closely conserved along the chromosome in different human populations. It is therefore vital to take this into account when studying recombination and searching for signatures of natural selection.The power spectrum showed the proportion of variance is highest in the fine-mid scales, which is expected given the existence of hotspots and the distribution of recombination across the chromosomes. This is evidenced further by the correlation between detail coefficients increasing with scale, thus most differences between the populations are at the fine scale. Coherence analysis showed that differences are not uniform across the chromosome, at any scale. Whilst it should be noted that our analysis is limited to a subset of a single chromosome, and some variation between chromosomes would be expected, the principles highlighted will apply more broadly, even if there is regional variation

The results reported here are broadly in line with other results and expectations. The proportion of variance at each scale for all datasets is very similar to that reported by Bherer *et al.* [47] where male and female recombination rates were compared and similar conclusions were drawn. The comparison between two *D. melanogaster* populations in Chan *et al*. [30] shows a similar increase in correlation between the populations as the scale size increases, as in Figure 2. This shows that the shape of correlations between human populations are similar in other species where there is a geographic distance between populations.

It is somewhat surprising that the African recombination datasets were more correlated to each other than the European datasets were, over every scale. It is known that there is far more genetic diversity in Africa than there is in Europe [63]. The higher correlation could be because the Baganda and Zulu populations are more closely related than the two European datasets: they both speak languages from the Bantoid family, a subgroup of the Niger-Congo languages [51], although it should be noted that this is a broad classification, encompassing hundreds of languages and dialects spoken by hundreds of millions of people in present sub-Saharan Africa [64]. Other research using LD-maps has shown the two populations are much more similar to each other than to Ethiopians [65]. The so-called “Bantu expansion”, where West African Bantu-speaking people spread west and south across the continent, happened around 5,600 years ago, integrating with the people who had been there for much longer [64]. Therefore, while their languages may have a fairly recent common ancestor, an estimate of the split between the Baganda and Zulu populations is around 29,000 years ago [66], so this is unlikely to be the cause. Another theory could be that the European pairing has lower correlations due to selection. The ‘Out of Africa’ event and subsequent spreading across the Eurasian continent engendered huge selective pressures as new environments, diets, and climates were encountered [67]. Recombination hotspots will move around the genome due to selective pressure in locations where it would be beneficial for alleles along a chromosome to be separated [68, 69]. However, the African populations will have experienced their own set of selective pressures and so this may not explain the difference. Regardless of the reasons for the difference, it has been shown there is a difference between African and European recombination rates. Therefore, it is important when using recombination maps for analysis, for example when searching for regions under selection, to use maps made for the specific population being analysed. This is especially true when analysing genetic data on the fine scale [34].

Previous work has been undertaken to describe the differences between recombination rates of human populations, although none appear to have used wavelet methods. The work of Hinch *et al*. [33] on African American recombination maps reinforces the conclusions drawn here regarding similarity on the very wide scale, and differences on at the very fine scale. They conclude that variations in PRDM9 are the driving force behind the differences in hotspot location. Looking further back into the evolution of *Homo sapiens*, Lesecque *et al*. [70] consider the recombination differences between modern humans and Denisovans, whose lineages are thought to have split around 750,000 years ago [71], and discovered that they share few hotspots, even though the motif associated with hotspots in modern humans was already being actively targeted by PRDM9 before the split.

In conclusion, we have highlighted the potential for wavelets to support the interrogation of recombination (and LD-based estimates thereof), allowing multiple resolutions to be considered in parallel as opposed to arbitrary size cut-offs. Furthermore, variability between populations was shown, highlighting the need for appropriate population genomic data to be used for LD-based linkage map generation.

## ACKNOWLEDGEMENTS

This work was supported by funding from the University of Southampton Institute for Life Sciences, Faculty of Medicine and Department of Mathematics. The authors acknowledge the use of the IRIDIS High Performance Computing Facility, and associated support services at the University of Southampton, in the completion of this work.

## SUPPLEMENTARY LEGENDS

**Supplementary Figure 1. Principal component analysis (PCA) of Wellderly individuals.**

PCA was used to identify a homogenous population sample dataset using PLINK (v.1.90beta) [52], utilising genotypes pruned using a window size of 50 variants, an LD (r^2^) value of 0.5 and shifting by 5 variants each step. The data were then clustered into five clusters using a k-means clustering algorithm in R (v3.3.0) [53], using the Hartigan-Wong algorithm in the stats package (v3.4.1) [72] with starting sets equal to 25 and maximum iterations set to 1000, using the first four components. Individuals in the largest cluster were considered to be of European ancestry based on the predominant self-reported ancestry of the cohort and PCA analysis alongside CEU samples, and all other individuals excluded from further analyses.

**Supplementary Figure 2. Comparison of Bherer and LDhat recombination maps**

The top figure shows the refined European sex-averaged linkage map created by Bherer [47](in blue) and the estimated recombination rate map generated by LDhat (in solid black), for a 31 Mb region of chromosome 22. The Bherer map is in centimorgans (cM) and LDhat in historical centimorgans. The lower figure shows the same data transformed into centimorgans per megabase (cM/Mb).

**Supplementary Figure 3. Scatterplots of LDhat against Bherer map**

The top figure shows the LDhat map against the refined European sex-averaged linkage map created by Bherer (both 1 kb bins). The blue line is a simple linear regression fitted using R. The R squared value and p-value are reported. The lower figure is scatterplot of the log_10_ data. A Kendall’s Tau value of 0.41 was calculated (p-value < 2.2×10^-16^), indicating a non-zero relationship between the two maps.

**Supplementary Figure 4. European cold spot**

The data for the four datasets in log10 cM/Mb along chromosome 22. This figure shows a cold spot between 32.3 Mb and 32.55 Mb in the European (CEU and Wellderly) data, but not in the African (Bganda and Zulu) data.

