## Supplementary material for "Wavelet analysis of human recombination rates demonstrates divergence on fine scales"

### Supplementary FIGURES

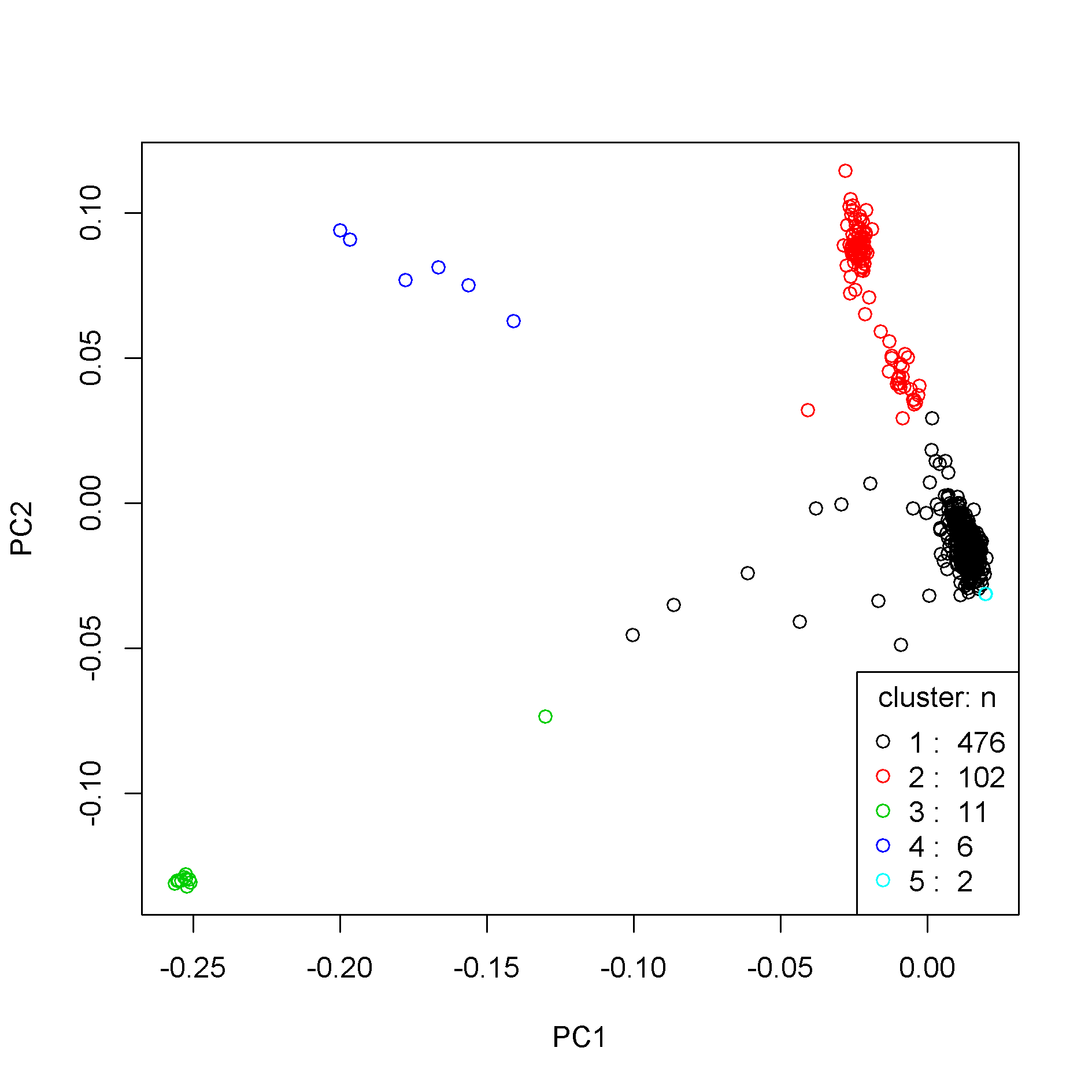

Supplementary Figure 1. Principal component analysis (PCA) of Wellderly individuals. PCA was used to identify a homogenous population sample dataset using PLINK (v.1.90beta) [1], utilising genotypes pruned using a window size of 50 variants, an LD (*r^2^*) value of 0.5 and shifting by 5 variants each step. The data were then clustered into five clusters using a k-means clustering algorithm in R (v3.3.0) [2], using the Hartigan-Wong algorithm in the stats package (v3.4.1) [3] with starting sets equal to 25 and maximum iterations set to 1000, using the first four components. Individuals in the largest cluster were considered to be of European ancestry based on the predominant self-reported ancestry of the cohort and PCA analysis alongside CEU samples, and all other individuals excluded from further analyses.

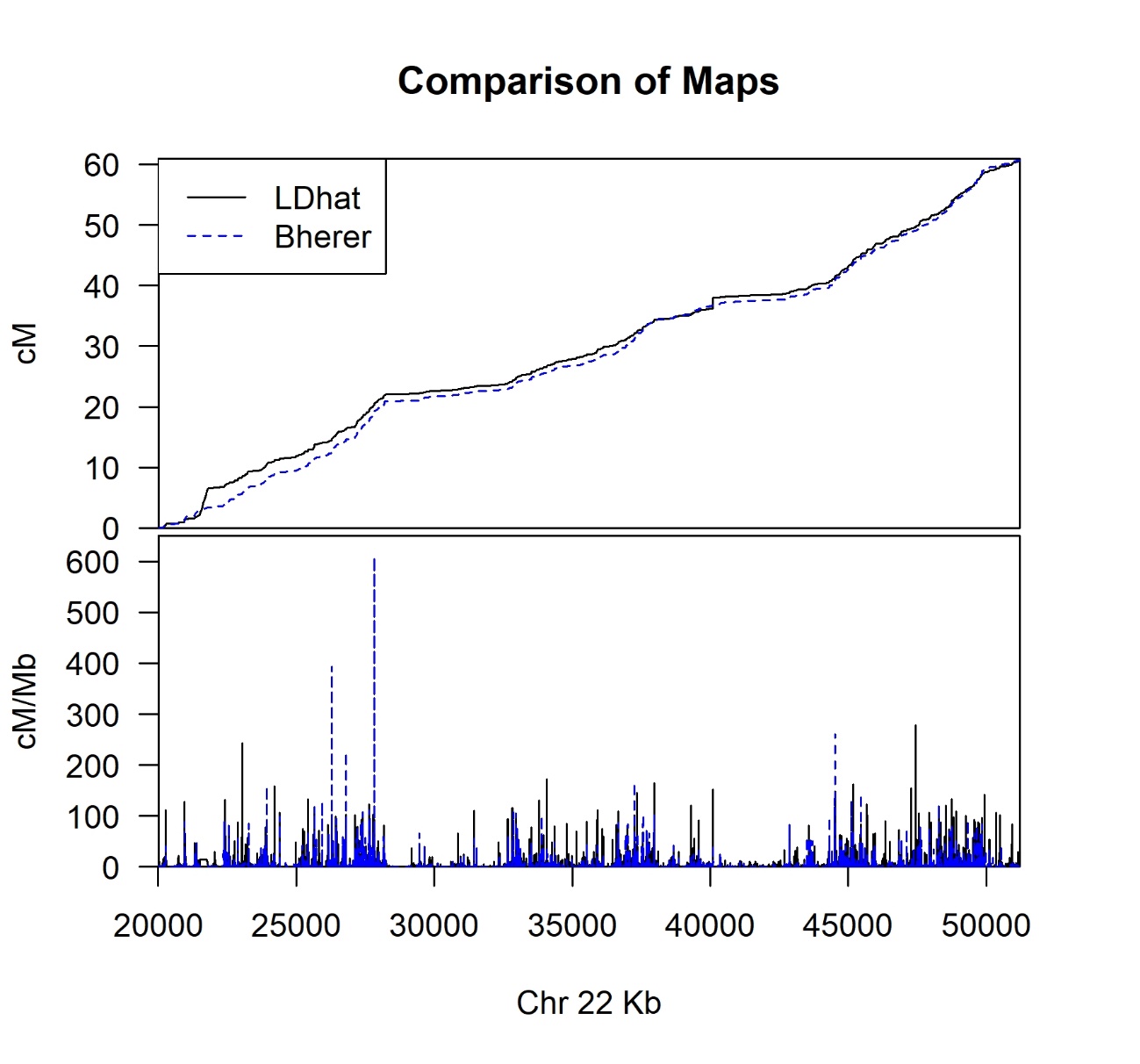

Supplementary Figure 2. Comparison of Bherer and LDhat recombination maps

The top figure shows the refined European sex-averaged linkage map created by Bherer [4](in blue) and the estimated recombination rate map generated by LDhat (in solid black), for a 31 Mb region of chromosome 22. The Bherer map is in centimorgans (cM) and LDhat in historical centimorgans. The lower figure shows the same data transformed into centimorgans per megabase (cM/Mb).

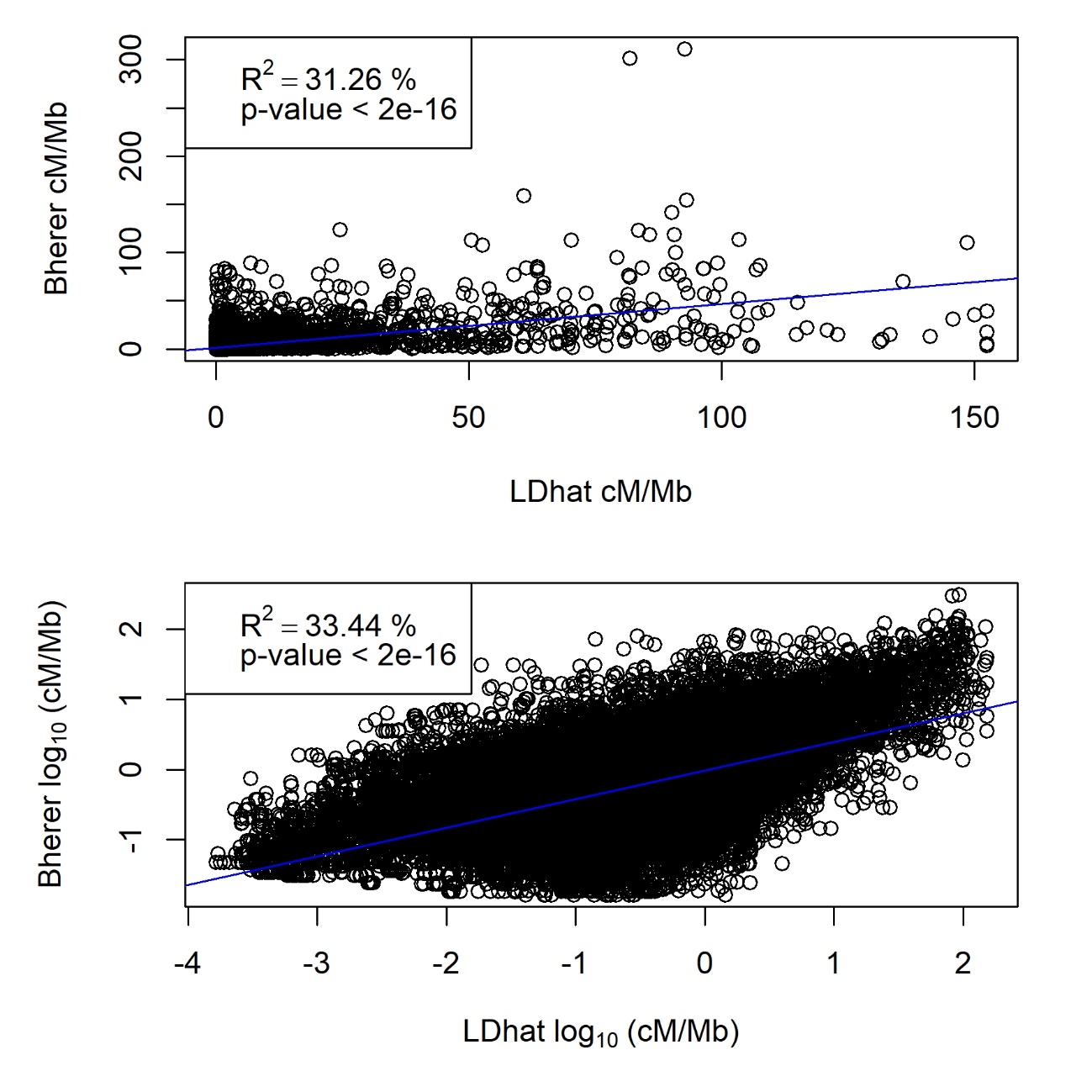

Supplementary Figure 3. Scatterplots of LDhat against Bherer map

The top figure shows the LDhat map against the refined European sex-averaged linkage map created by Bherer (both 1 kb bins). The blue line is a simple linear regression fitted using R. The R squared value and p-value are reported. The lower figure is scatterplot of the log_10_ data. A Kendall’s Tau value of 0.41 was calculated (p-value < 2.2x10^-16^), indicating a non-zero relationship between the two maps.

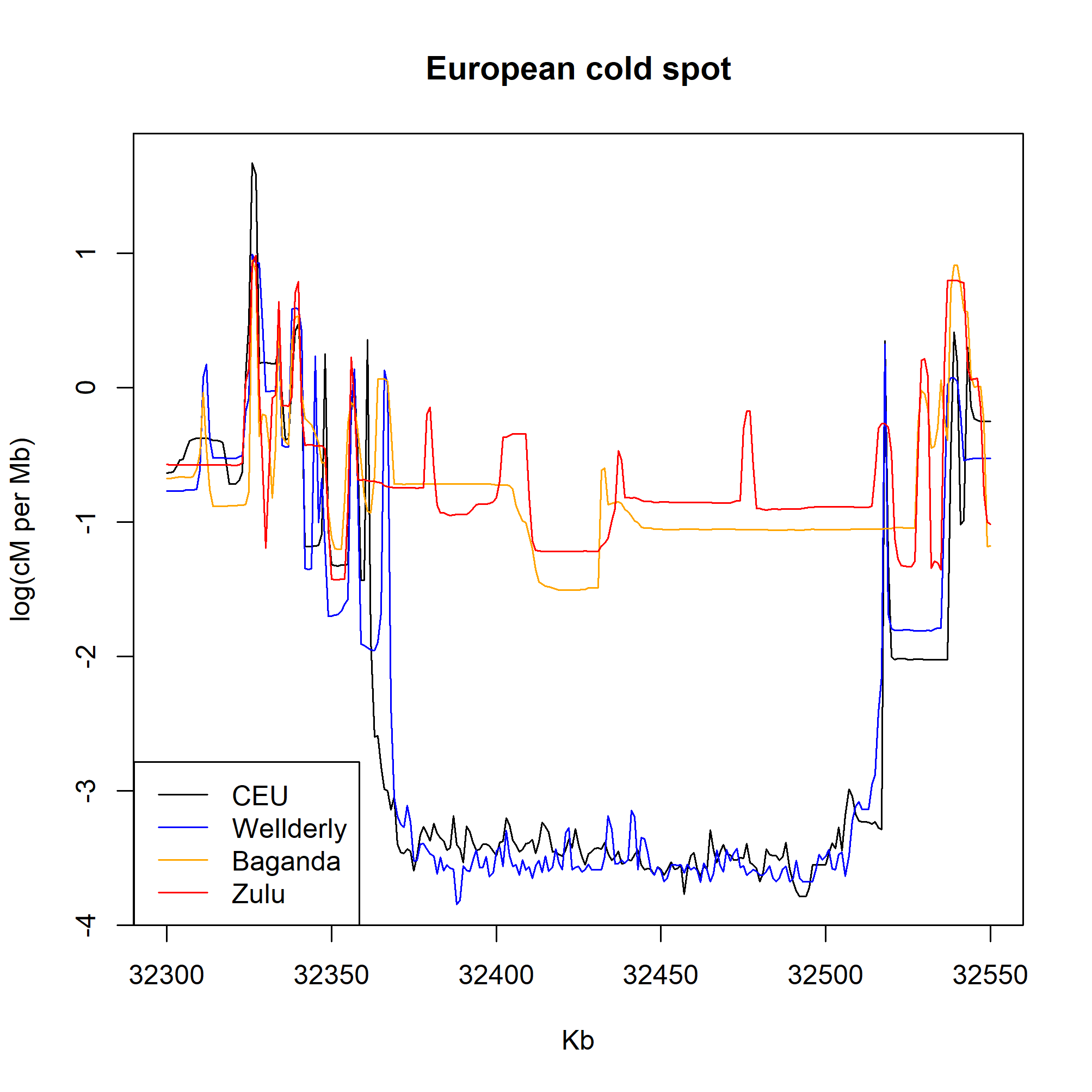

Supplementary Figure 4. European cold spot

The data for the four datasets in log10 cM/Mb along chromosome 22. This figure shows a cold spot between 32.3 Mb and 32.55 Mb in the European (CEU and Wellderly) data, but not in the African (Bganda and Zulu) data.
